# Alphaherpesvirus pUL21 homologues use non-canonical motifs to compete with cellular adaptors for protein phosphatase 1 binding

**DOI:** 10.1101/2025.07.22.666160

**Authors:** Holly Monkhouse, Daniela S. Carter-Lopez, Tomasz H. Benedyk, Janet E. Deane, Stephen C. Graham

## Abstract

Protein phosphatase 1 (PP1) is a key regulator of cellular phosphorylation and its activity is regulated via binding to cellular regulatory proteins via conserved short linear motifs (SLiMs). The herpes simplex virus (HSV)-1 protein pUL21 binds PP1 via the TROPPO motif, which lacks sequence similarity to canonical PP1-binding SLiMs. Here, we combine structure prediction, mutagenesis, and biophysical assays to elucidate the molecular basis of this interaction. AlphaFold2-Multimer structural models suggest that the TROPPO motifs of pUL21 and of pORF38, the varicella-zoster virus homologue of pUL21, bind the same hydrophobic groove on PP1 as RVxF and ϕϕ[xF] motifs, forming an extended β-sheet that bridges PP1 and the N-terminal domain of pUL21 or pORF38. Site-directed mutagenesis of both pUL21 and PP1 confirms key predicted interactions. Competition fluorescence polarisation confirms that pUL21 competes directly with cellular PP1 RvXF motifs for PP1 binding, albeit with lower affinity. Substituting key residues of the pUL21 TROPPO motif to resemble a canonical RVxF sequence increases PP1 binding, suggesting that alphaherpesviruses have evolved suboptimal motifs to fine-tune phosphatase recruitment that may balance viral kinase and phosphatase activities during infection. Our findings reveal a novel mechanism of PP1 recruitment by viral proteins and suggest that many other PP1 regulators may possess non-canonical binding motifs and thus remain undiscovered.

## Introduction

Phosphorylation is critical to the control of most cellular processes. Viruses have learnt to exploit this, encoding kinases that regulate cell cycle progression, modify gene expression, overcome innate immune restriction and prevent apoptosis (1, 2). Similarly, multiple viruses modulate the dephosphorylation of both cellular and viral target proteins within infected cells. Herpes simplex virus (HSV)-1 protein ICP34.5 (a.k.a. γ134.5) counteracts the antiviral innate immune response by stimulating dephosphorylation of eIF2α to relieve host transcriptional shutdown (3, 4). The HIV Tat protein interacts with the cellular enzyme protein phosphatase 1 (PP1) to promote viral gene transcription (5, 6). Additionally, the measles virus V protein suppresses innate immune signalling by sequestering PP1, thereby preventing dephosphorylation of the cytosolic RNA sensor MDA5 that is required for its activation (7). Given the multitude of cellular signalling pathways that are regulated by protein phosphorylation, interfering with virus-mediated protein dephosphorylation represents a promising new avenue for the development of potent antiviral therapies (6, 8).

PP1 is a highly abundant cellular serine/threonine phosphatase. PP1 has low intrinsic specificity and it recruits its substrates via regulatory proteins (9, 10), also known as adaptor proteins, PP1 Interacting Proteins (PIPs) or Regulators of Protein Phosphatase One (RIPPOs) (11). These regulatory proteins utilise small linear motifs (SLiMs) to bind different surfaces of PP1 (9, 10). The most abundant and well characterised of these is the RVxF motif, which binds to a hydrophobic groove on the surface of PP1. Over 90% of cellular PP1 adaptors have an RVxF motif (12), which has the consensus sequence [KR][KR][VI]ψ[FW], where ψ is any amino acid except FIMYDP (13). Many PP1 regulatory proteins possess more than one PP1-interacting SLiM (reviewed in (10)) allowing combinatorial control of PP1 binding (14). One example of an ‘accessory’ SLiM is the ϕϕ[xF] motif, first identified via structural characterisation of PP1 in complex with spinophilin (15), which lies approximately 6–8 residues downstream of the RVxF motif and is present alongside the RVxF motif in most PP1 regulatory proteins (16). Like their cellular counterparts, viral PP1 regulatory proteins commonly contain an RVxF motif that is critical for PP1 binding (5, 7, 17), but the presence of additional SLIMs in these adaptors is less well explored.

HSV-1 pUL21 was recently identified as a novel viral PP1 regulatory protein that promotes virus replication and spread. HSV-1 pUL21 and its homologues across the alphaherpesviruses comprise two folded domains (18, 19) that are linked by a flexible linker domain (20). PP1 binds pUL21 via a novel short linear motif, termed the Twenty-one Recruitment Of Protein Phosphatase One (TROPPO) motif, that is present in the linker region and is conserved across alphaherpesviruses. Mutation of the TROPPO motif abolishes the ability of either HSV-1 pUL21 or the Varicella-Zoster virus (VZV) homologue (pORF38) to bind PP1. Loss of PP1 binding results in hyperphosphorylation of multiple proteins, both viral and cellular, in HSV-1 infected cells. pUL21 binding to PP1 regulates phosphorylation of components of the viral nuclear egress complex (20), which remodels the nuclear membrane to support virus maturation (21, 22). The pUL21:PP1 interaction also leads to dephosphorylation and thus hyperactivation of the ceramide transport protein CERT, altering sphingolipid metabolism in infected cells (23). This TROPPO motif does not bear obvious relation to known PP1 interacting motifs and pUL21 does not have any other known PP1 binding motifs such as an RVxF motif. The molecular basis of pUL21 binding to PP1 was thus unclear.

While inhibition of the ubiquitous and highly active enzyme PP1 is toxic, disrupting specific PP1 activities via blocking of specific interactions is a promising avenue for the design of new therapies (24). The lack of homology with ‘RVxF’ motifs used by cellular PP1-binders suggested that the viral TROPPO motif might bind a novel surface on PP1 and thus could be specifically targeted by compounds that would inhibit virus replication and spread with minimal side-effects, as cellular functions of PP1 would remain uninhibited. We thus sought to determine how the TROPPO motif of HSV-1 pUL21 and its homologues bind PP1.

## Methods

### Plasmids

Bacterial expression plasmids encoding the mouse PP1γ catalytic domain (residues 7–300)-H_6_ and pUL21-H_6_, and mammalian expression vectors encoding pUL21(FV242AA)-GFP, pORF38-GFP and pORF38(FV255AA)-GFP, was described previously (20). We note that the mouse and human PP1γ catalytic domains share 100% amino acid identity. pORF38-NLT-H_6_ was generated by sub-cloning residues 1–263 of ORF38 into the vector pOPTnH (25). pUL21-RVxF-H_6_ was generated by inserting the mutations A237V and V239F into pUL21-H_6_ by QuikChange site-directed mutagenesis. Human PP1α was cloned from HeLa cell cDNA into a vector derived from pF5K (Promega) with an N-terminal Myc epitope tag via restriction-based cloning, and point mutations were introduced into Myc-PP1α by QuikChange mutagenesis. A plasmid encoding pUL21-GFP was generated by sub-cloning a codon-optimised synthetic UL21 gene (GeneArt) into pEGFP-N1 (Clontech), yielding an identical pUL21-GFP amino acid sequence as used in (20). pUL21-ΔL, -ΔR and -ΔLR were generated from this vector via inverse PCR, and point mutations were introduced via QuikChange mutagenesis.

### Peptides

Peptides were commercially synthesised to >95% purity (GenScript) as follows: GADD34 RVxF+ϕϕ[xF] (residues 552–568) with fluorescein isothiocyanate (FITC) attached via an N-terminal aminohexanoic (Ahx) linker (sequence: [FITC-Ahx]-KARKVRFSEKVTVHFLA); pUL21 TROPPO (residues 234–250, sequence: AKRATVSEFVQVKHIDR); pOPF38 TROPPO (residues 246–263; sequence: KSDHITLSNFVQIRTIPR). TROPPO motif peptides were dissolved in aqueous buffers and GADD34 RVxF+ϕϕ[xF] was dissolved in dimethyl sulfoxide (DMSO).

### Recombinant protein purification

Proteins were expressed using *Escherichia coli* T7 Express *lysY/I^q^* cells (New England Biolabs) grown in 2×TY medium at 37°C to OD_600_ 0.8–1.2 before cooling to 22°C and induction of protein expression using 0.4 mM IPTG. Cells were harvested at 16–20 hours post-induction and pellets stored at −70°C until required. Cells were resuspended in chilled lysis buffer (20 mM Tris pH 7.5, 20 mM imidazole, 500 mM NaCl, 1.4 mM β-mercaptoethanol 0.5 mM MgCl_2_ and 0.05% TWEEN-20 [plus 1 mM MnCl_2_ for PP1]) that was supplemented with 200 µL EDTA-free protease inhibitor cocktail (Merck) and 400 U bovine DNase I (Merck). Cells were lysed using a TS series cell disruptor (Constant Systems) at 24 kpsi and lysates were clarified (40,000×g, 30min, 4 °C) before incubation with Ni-NTA agarose (Qiagen) for 1 h at 4°C. The resin was washed with ≥20 column volumes of wash buffer (20 mM Tris, 20 mM imidazole and 500 mM NaCl [plus 1 mM MnCl_2_ for PP1]) at pH 7.5 (PP1γ and pORF38-NLT) or pH 8.5 (pUL21 and pUL21-RVxF). Protein was eluted using wash buffer supplemented with additional imidazole (final concentration 250 mM) and proteins were subjected to size-exclusions chromatography using HiLoad 16/600 Superdex 75 (PP1) or Superdex 200 (other) columns equilibrated in 20 mM Tris pH 8.5, 500 mM NaCl, 1 mM DTT (for pUL21 and pUL21-RVxF) or 50 mM HEPES pH 8.0, 500 mM NaCl, 0.5 mM TCEP (for PP1 and pORF38-NLT). Fractions were analysed by SDS-PAGE and those containing the desired protein were pooled, concentrated using centrifugal concentrators (Millipore) and stored at 4°C (short-term) or snap-frozen in liquid nitrogen for long-term storage at −70°C.

### Isothermal Titration Calorimetry (ITC)

ITC experiments were performed using a MicroCal PEAQ-ITC automated calorimeter (Malvern Panalytical). The solvent (ITC buffer) was 50 mM HEPES pH 8.0, 500 mM NaCl, 0.5 mM TCEP. Lyophilised peptides were dissolved into ITC buffer and their concentration was estimated from dry mass. Proteins were diluted in ITC buffer and their concentration was estimated from the theoretical extinction coefficient at 280 nm (26). Titrations were conducted at 25°C using 12 x 3 µL injections with syringe and cell contents as listed in Table S1. For all reagents, control titrations (syringe into buffer or buffer into cell) showed very low non-specific heat evolution. Data were analysed using MicroCal PEAQ-ITC analysis software (Malvern Panalytical) and fitted using a one-site binding model.

### Fluorescence polarisation anisotropy

Fluorescence anisotropy was measured using a SpectraMax i3 microplate reader (Molecular Devices) with wavelengths 485 ± 20 nm (excitation) and 535 ± 25 nm (emission). Samples (100 µL per well) were dispensed into low-bind black half-area 96-well microtitre plates (Corning) and fluorescence anisotropy was recorded at 27°C. Experiments were performed in 50 mM HEPES pH 8.0, 500 mM NaCl, 0.05% TWEEN-20 unless otherwise noted. For all titrations, fluorescent GADD34 RVxF+ϕϕ[xF] peptide was diluted from a 5 µM stock in DMSO into buffer for a final concentration of 4 nM (0.08% DMSO). To measure association, 2 nM peptide was incubated with a serial dilution of PP1γ catalytic domain. For competition assays a serial dilution of the competitor was mixed 1:1 with a pre-formed complex of 4 nM GADD34 peptide and 500 nM PP1γ catalytic domain, to yield a final concentrations of 2 nM GADD34 peptide, 250 nM PP1γ and competitor as indicated. The PP1γ dissociation constant (*K*_D_) was calculated by fitting the grating factor corrected anisotropy data to a one-site binding equilibrium model in Prism version 7 (GraphPad). For competition experiments, IC50 values were calculated by fitting the grating factor corrected anisotropy data to a four-parameter inhibitor concentration response curve using Prism version 7 (GraphPad).

### Mammalian cell culture

Mycoplasma-free, human embryonic kidney 293 T (HEK 293T) cells (American Type Culture Collection #CRL-3216) were maintained in Dulbecco’s modified Eagle’s medium (DMEM) with high glucose (Merck), supplemented with 10% (v/v) heat-inactivated fetal calf serum and 2 mM _L_-glutamine (complete DMEM) in a humidified 5% CO_2_ atmosphere at 37 °C.

### Immunoprecipitation of GFP-tagged bait protein from transfected cells

Monolayers of HEK 293T cells were transfected with 7.7 μg of DNA per 9 cm dish, using 18 μg/mL 25 kDa branched polyethylenimine (Merck) or using TransIt-LT1 (Mirus) in accordance with the manufacturer’s instructions. Cells were harvested 24 h post-transfection and incubated for 30 min in ice-cold 10 mM Tris pH 7.5, 150 mM NaCl, 0.5 mM EDTA, 0.5% Igepal CA-630 (a.k.a. NP-40), 1% (v/v) EDTA-free protease inhibitor cocktail (Merck) and incubated (4°C, 30 min) before clarification at 21,000×g for 10 min at 4 °C. Lysate protein concentrations were measured using the bicinchoninic acid (BCA) assay (Pierce) and normalised before affinity capture using GFP-Trap resin (ChromoTek) following the manufacturer’s protocol. Samples were eluted by heating the resin to 95°C for 5 min in 45 μL 2×SDS-PAGE loading buffer. Proteins were separated by SDS-PAGE and transferred to 0.45 μm nitrocellulose membranes (PerkinElmer or Cytiva). Membranes were stained with Ponceau S and imaged using a G:Box XX9 (Syngene) before blocking in Tris-buffered saline with 0.1% TWEEN-20 (TBS-T) supplemented with 5% (w/v) non-fat milk powder. Immunoblotting was performed using antibodies diluted in blocking buffer as listed below and immunoblots were imaged using an Odyssey CLx (LI-COR Biosciences). Densitometry was performed using Image Studio Lite version 5.2 (LI-COR Biosciences) and statistical tests were performed using Prism version 7 (GraphPad).

### Antibodies

The following primary antibodies and dilutions were used for immunoblotting: rabbit anti-CERT 1:10,000 (Abcam #Ab72536), mouse anti-PP1α 1:1000 (Santa Cruz #sc-271762), mouse anti-Myc 1:4000 (Millipore #05-724), rat anti-tubulin (clone YL1/2) hybridoma supernatant 1:40 (27). Secondary antibodies from LI-COR Biosciences were diluted 1:10,000 as follows: IRDye 680RD goat anti-rat (#926-68029) and goat anti-mouse (#926-68020), or IRDye 800CW goat anti-rabbit (#926-32221).

### Structure prediction and analysis

Structures were predicted using a locally installed version of ColabFold version 1.5.3. Input sequences were human PP1γ catalytic domain (residues 7-300, UniProt P36873) plus either HSV-1 strain KOS pUL21 (residues 1-535, UniProt ID: F8RG07) or VZV strain Dumas pORF38 (residues 1-541, Uniprot P09289) in complex with PP1. Mean lipophilicity maps were calculated using ChimeraX (28) and molecular graphics were generated using an open-source build of PyMOL (Schrödinger).

## Results

### The TROPPO motif alone is insufficient to bind PP1

Prior studies of cellular regulatory proteins showed that peptides containing the RXvF and ϕϕ[xF] motifs are sufficient to bind the catalytic domain of PP1 (29). Using fluorescence anisotropy, we confirmed that a peptide containing the RVxF and ϕϕ[xF] motifs of GADD34 (a.k.a. PP1 regulatory subunit 15A) binds purified PP1γ catalytic domain (residues 7–300) with 144 ± 84 nM affinity (mean ± SEM, n = three independent experiments each performed in technical triplicate) (Fig. 1A), similar to previous isothermal titration calorimetry (ITC) affinity measurements for the same peptide sequence (29). Competition fluorescence anisotropy experiments were performed, whereby peptides containing the TROPPO motifs of pUL21 or pORF38 were titrated into a pre-formed complex of PP1 plus the fluorescent GADD34 peptide. Neither TROPPO-containing peptide was able to displace the GADD34 peptide, even when present at over 1000-fold molar excess (Fig. 1B). This suggested that either the TROPPO motifs do not bind PP1, or they bind a different site on the PP1 surface that does not overlap with the RVxF binding groove. ITC was employed to distinguish between these possibilities. Surprisingly, titrating high concentrations (500–1000 µM) of pUL21 or pORF38 TROPPO-containing peptides against the PP1 catalytic domain did not evolve heats above background levels (Fig. 1C,D, Table S1), suggesting that neither peptide is sufficient to bind PP1.

**Figure 1.**
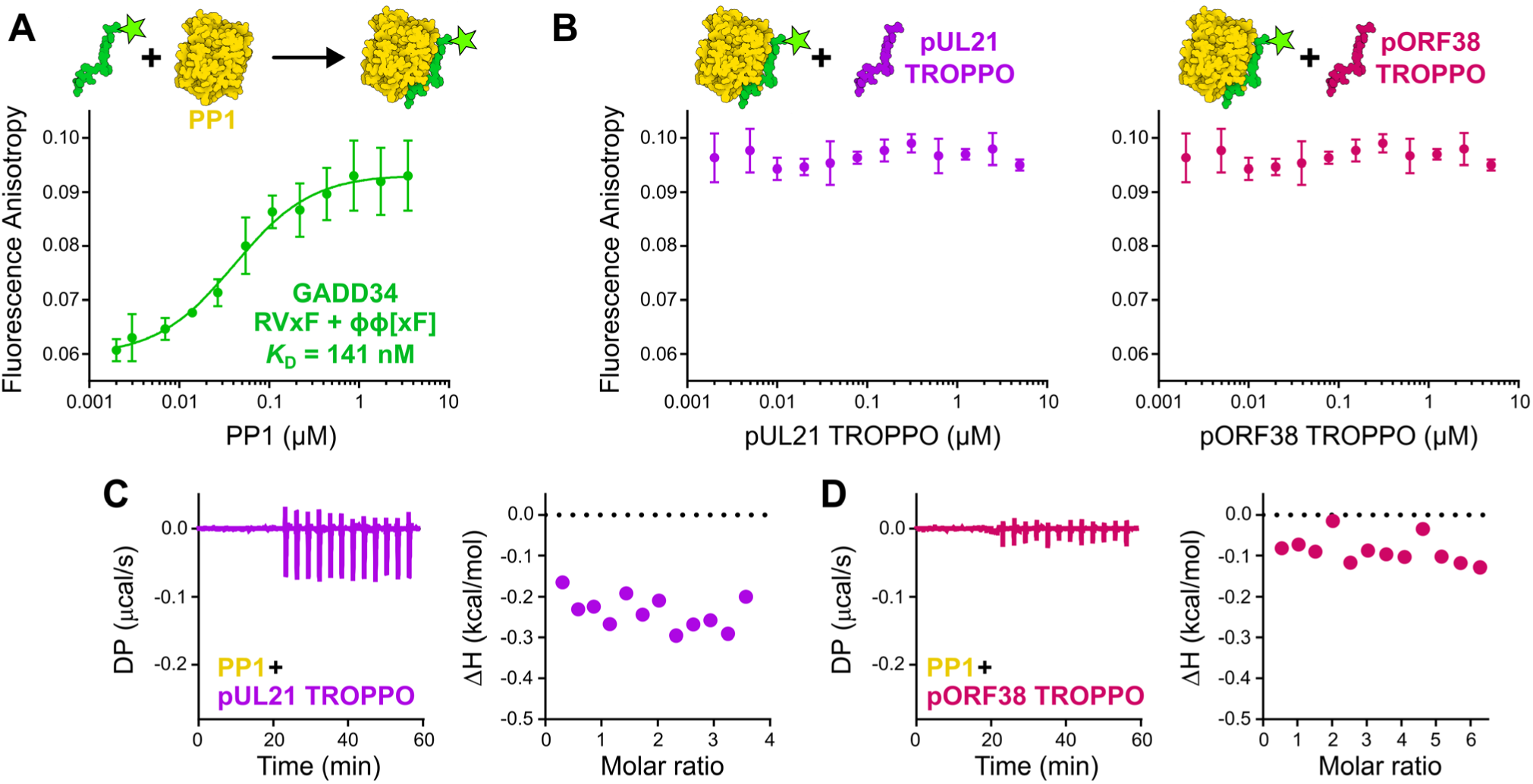
Alphaherpesvirus TROPPO peptides do not bind PP1. **A** Fluorescence anisotropy of PP1γ catalytic domain (residues 7–300) binding 2 nM fluorescently labelled peptide containing the GADD34 RVxF and ϕϕ[xF] motifs (residues 552–568). Mean ± SD is shown for technical triplicate measurements. Affinity (*K*_D_) is average of three independent experiments. **B** Competition fluorescence anisotropy of peptides containing the alphaperhesvirus TROPPO motif. Pre-formed complexes of 2 nM fluorescent GADD34 peptide with 250 nM PP1 were incubated with increasing concentrations of TROPPO-containing peptides from pUL21 (residues 234–250; left) and pORF38 (residues 246–263; right). Neither peptide successfully displaced GADD34 RVxF. Mean ± SD is shown for technical triplicate measurements. **C, D** Isothermal titration calorimetry of PP1γ catalytic domain with TROPPO containing peptides from **C** pUL21 and **D** pORF38. For each, baseline-corrected differential power (DP) versus time (left) and integrated change in enthalpy (ΔH) versus molar ratio (right) are shown. Figures are representative of three (**C**) or two (**D**) independent experiments.

### The C-terminal region of the pUL21 linker is dispensable for PP1 binding

The N-terminal domain and linker region of pUL21 are necessary for immunoprecipitation (IP) of PP1, with mutation of the TROPPO motif within the linker domain efficiently disrupting this binding (20). While the TROPPO motif is absolutely conserved across alphaherpesvirus pUL21 homologues, the remainder of the pUL21 linker regions are only poorly conserved (Fig. 2A). Strikingly, the region of the linker between the TROPPO motif and the C-terminal domain is significantly longer in HSV-1 and HSV-2 pUL21 than in other herpesviruses such as VZV, the pUL21 homologue of which (pORF38) is also capable of binding PP1 (20).

**Figure 2.**
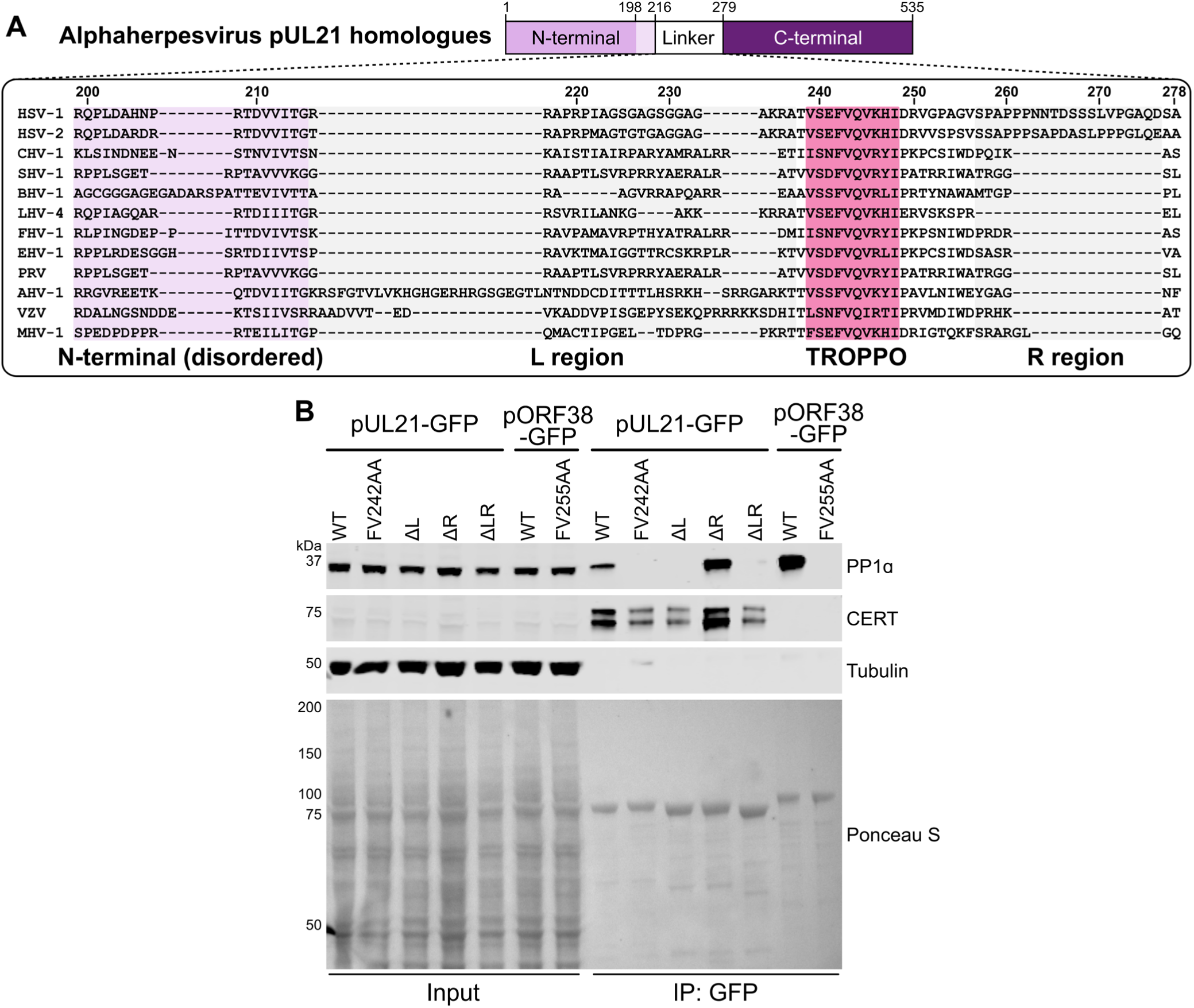
Contribution of the pUL21 linker region to PP1 binding. **A** Alignment of alphaherpesvirus pUL21 homologue sequences spanning the linker between pUL21 N- and C-terminal domains, plus the N-terminal domain residues that were disordered in the pUL21-N crystal structure (18). The following sequences were aligned using ClustalW (Abbreviation and Uniprot ID are shown in parentheses): HSV-1 (HSV1, P10205), HSV-2 (HSV2, G9I242), cercopithecine herpesvirus 2 (CHV-2, Q5Y0T2), saimiriine herpesvirus 1 (SHV-1, E2IUE9), bovine alphaherpesvirus 1 (BHV-1, Q65563), leporid alphaherpesvirus 4 (LHV-4, J9QYM9), feline herpesvirus 1 (FHV-1, D1FXW1), equine herpesvirus 1 (EHV-1, P28972), pseudorabies virus (PRV, Q04532), anatid herpesvirus 1 (AHV-1, A4GRJ2), varicella-zoster virus (VZV, Q6QCT9), turkey herpesvirus (MHV-1, Q9DPR5). The regions of the linker preceding (L region, residues 216–237, grey) or following (R region, residues 257–276, grey) the TROPPO motif (pink) are highlighted. **B** HEK293T cells were transfected with plasmids expressing wild-type (WT) or mutated HSV-1 pUL21-GFP or VZV pORF38-GFP. At 24 hours post-transfection the cells were lysed, subjected to immunoprecipitation using a GFP affinity resin, and captured proteins were subjected to SDS-PAGE and immunoblotting using the listed antibodies. Ponceau S (PonS) staining of the nitrocellulose membrane before blocking is shown, confirming efficient capture of GFP-tagged proteins. Figure is representative of two independent experiments.

To probe the contribution of the linker region outside the TROPPO motif to PP1 binding, deletion mutants of HSV-1 pUL21 were designed lacking linker residues that precede the TROPPO motif (residues 216–237; pUL21ΔL), linker residues that follow the TROPPO motif (residues 257–276; pUL21ΔR), or both flanking regions (pUL21ΔLR). Wild-type and mutant pUL21, and the pUL21 homologue ORF38 from VZV, were expressed as GFP fusion proteins and used for co-IP analysis following transient transfection in HEK293T cells (Fig. 2B). As previously reported, pUL21 and pORF38 efficiently precipitate endogenous PP1α and the interaction is disrupted by mutation of the TROPPO motif (pUL21-FV242AA and pORF38-FV255AA). Strikingly, removal of the N-terminal region of the pUL21 linker (pUL21ΔL or pUL21ΔLR) completely abolishes PP1α binding, whereas binding is retained when the C-terminal region of the pUL21 linker is removed (pUL21ΔR). All pUL21 constructs retain CERT binding, consistent with localisation of CERT-binding to the C-terminal domain of pUL21 (20, 23) and confirming the correct expression and folding of these mutants.

### Structural model of the pUL21:PP1 and pORF38:PP1 complexes

Extensive attempts to co-crystallise WT or ΔR pUL21 with the catalytic subunit of PP1 proved unsuccessful. Therefore, to investigate the relative contributions of the N-terminal domain, linker and TROPPO motif to PP1 binding, the structure of HSV-1 pUL21 (Fig. 3A) and VZV pORF38 (Fig. 3B) in complex with the catalytic domain of human PP1γ were predicted using AlphaFold2-Multimer (30) via the Colabfold pipeline (31). The models for pUL21 (Fig. 2A, pLDDT/ipTM = 86.0/0.936) and pORF38 (Fig. 2B, pLDDT/ipTM = 85.4/0.918) predict that the TROPPO motif and several segments of the linker region preceding it associate with the N-terminal domain and with PP1. The per-residue confidence of the prediction (pLDDT) is high for residues of the ordered pUL21 and pORF38 N- and C-terminal domains and for the PP1 catalytic domain, plus for residues of the linker region that are predicted to interact with these domains (Fig. 3C). Analysis of the Predicted Aligned Error (PAE) for both models confirms that the orientation of the N-terminal domain and linker region up to the TROPPO motif with respect to PP1 is confidently predicted, while the relative orientations of linker region following the TROPPO motif and the C-terminal domains are not predicted with confidence (Fig. 3D).

**Figure 3.**
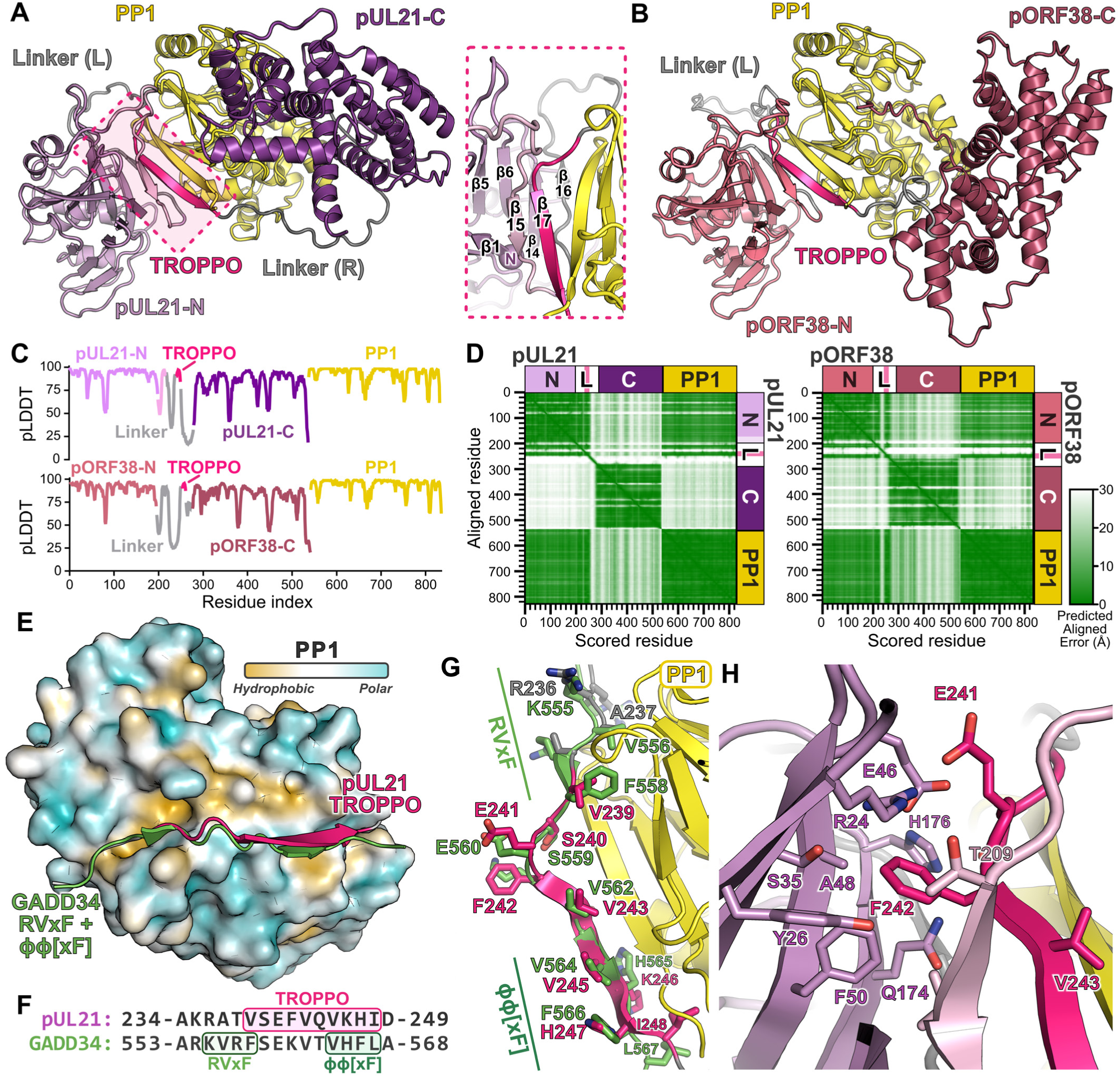
HSV-1 pUL21 and VZV pORF38 are predicted to bind the ‘RVxF’ hydrophobic groove of PP1. **A,B** AlphaFold2-Multimer models of human PP1γ catalytic domain (residues 7–300; yellow ribbons) in complex with **A** HSV-1 pUL21 (purple ribbons) or **B** VZV pORF38 (rose blush ribbons), with N- and C-terminal domains in lighter and darker shades, respectively. For each model, regions of the linker between the N- and C-terminal domains are coloured grey and the TROPPO motif is in bright pink. In **A** the region of the conserved N-terminal domain missing from the crystal structure (18) is light pink and *inset* shows the packing of predicted β-sheets at the pUL21-N:PP1 interface. **C** Per-residue predicted local distance difference test (pLDDT) of the pUL21:PP1 (top) and pORF38:PP1 (bottom) models, where values above 70 represent residues predicted with high local confidence. Domains are coloured as in (**A**, **B**). **D** Predicted aligned error of pUL21:PP1 (left) and pORF38:PP1 (right) models. Regions of the matrix where the relative orientations of residues are predicted with high confidence are shown in green. **E** Superposition of the pUL21:PP1 predicted structure on the crystal structure of PP1α in complex with the GADD34 RVxF+ϕϕ[xF] motif (PDB 4XPN) (29). PP1 is shown as a molecular surface coloured by mean lipophilicity, from orange (hydrophobic) to cyan (polar), and PP1-binding motifs are shown as ribbons. **F** Structure-based alignment of the pUL21 and GADD34 PP1-binding sequences, with SLiMs highlighted. **G** Predicted molecular interactions at the RVxF binding groove of PP1. Selected sticks are shown with carbon atoms green (GADD34) or coloured as in (**A**) (PP1 and pUL21). **H** Predicted interactions between pUL21-N and residues of the TROPPO motif. Selected sticks are shown with carbon atoms coloured as in (**A**).

Previously, small angle X-ray scattering showed that pUL21 comprises two ordered domains separated by a highly flexible linker (20). Crystallographic structures showed the N-terminal domain of pUL21 (pUL21-N) to possess a distinctive α/β fold, with two anti-parallel β-sheets that pack against surface-exposed α-helices (the ‘inner sheet’ and ‘lower outer sheet’) plus a third β-sheet that packs against the inner sheet (the ‘upper outer sheet’) (18). Despite pUL21 residues 1–216 being present in the crystallisation experiment and being conserved across alphaherpesviruses, only residues 1–198 were resolved in electron density. In the predicted complex, N-terminal domain residues 209–212 and residues 242–248 of the TROPPO motif are predicted to become ordered, forming two strands of anti-parallel beta sheet that serve as a bridge between the three-stranded pUL21-N upper outer sheet and the final β-sheet of the PP1 catalytic domain (Fig. 3A). Residues 221-223 of the Linker (L) region are also predicted to become ordered, forming a short β-strand that extends the pUL21-N inner sheet. We note that these newly-formed sheets are proximal to the amino terminus of pUL21, consistent with an inability of N-terminally GFP-tagged pUL21-N-plus-linker to bind PP1 (20).

All residues of the pUL21 TROPPO motif (_239_VSFVQVKHI_248_) are predicted to be in close association with PP1 (Fig. 3E). Surprisingly, the TROPPO motif is predicted to bind at the same surface hydrophobic groove as cellular RVxF-containing peptides, adopting a similar loop-plus-sheet conformation as GADD34 residues 553–568 (29). The first residue of the TROPPO and the three preceding (_236_RATV_239_) of pUL21 align structurally with residues _555_KVRF_558_ of the GADD34 RVxF motif (Fig. 3F), with pUL21 A237 and V239 predicted to bind the PP1 hydrophobic pockets occupied by GADD34 V556 and F558 (Fig. 3G). Most cellular regulatory proteins that bind PP1 possess a second PP1-binding SLiM, ϕϕ[xF], that is C-terminal to the RVxF (16). In the predicted complex, residues _244_VKHI_247_ of the pUL21 TROPPO adopt a similar conformation to the GADD34 ϕϕ[xF] residues _564_VHFL_567_ (Fig. 3G).

We previously showed that pUL21 mutations F242E or V243D are sufficient to abolish PP1 binding (23). The severity of the V243D mutation is readily explained by the complex prediction, this residue is predicted to interact with hydrophobic residues on the surface of PP1 (Fig. S1) and there is a valine at the equivalent position in GADD34 (Fig. 3G). The severity of the F242E mutation is less obvious when considering only the TROPPO motif, as the side chain is predicted to point away from PP1 (Fig. 3G). However, it becomes clearer when one considers the context of pUL21-N and the residues of the linker that are predicted to become ordered. The side chain of F242 is predicted to be largely buried, interacting with hydrophobic side chains of pUL21 A48 and F50, plus the hydrophobic face of the R24 guanidyl group (Fig. 3H). Mutation to a charged residue would be predicted to disrupt this packing. The TROPPO motif of pORF38 forms very similar interactions with both the pORF38 N-terminal domain and PP1 in the model of the pORF38:PP1 complex (Fig. S1). Loss of hydrophobic interactions would thus similarly explain why the FV255AA mutations abolished binding to PP1 (Fig. 2B) (20).

### Site-directed mutagenesis confirms the predictive power of the pUL21:PP1 model

While the models of pUL21 and pORF38 in complex with PP1 were compellingly consistent with both each other and with previous results, structurally informed mutagenesis is required to rigorously test their predictive power. To this end, single amino acids mutations were introduced in both PP1 and pUL21, and their ability to interact in cells was probed by coIP.

For PP1, we utilised three published mutations that have all been show to severely decrease PP1 regulatory protein binding: F257A (32), which is at the RVxF hydrophobic pocket into which the side chain of pUL21 V239 is predicted to bind (Fig. 4A); and L289R or C291R (33), which are on the final β-sheet of the PP1 catalytic domain and contribute to the RVxF hydrophobic surface with which pUL21 A237 is predicted to associate (Fig. 4A). HEK293T cells were co-transfected with pUL21-GFP and WT or mutant Myc-PP1, and lysates were subjected to GFP affinity capture before SDS-PAGE and immunoblotting. The coIP of all three mutants was profoundly less efficient than for WT Myc-PP1 (Fig. 4B). However, we consistently observed lower abundance of these mutants in the cell lysates used for IP analysis, suggesting lower expression or increased turnover in transfected cells. To account for this, the amount of input and bound Myc-PP1 was quantified and the ratio of bound:input was calculated (Fig. 4C), confirming that all three mutations significantly disrupt binding to pUL21-GFP.

**Figure 4.**
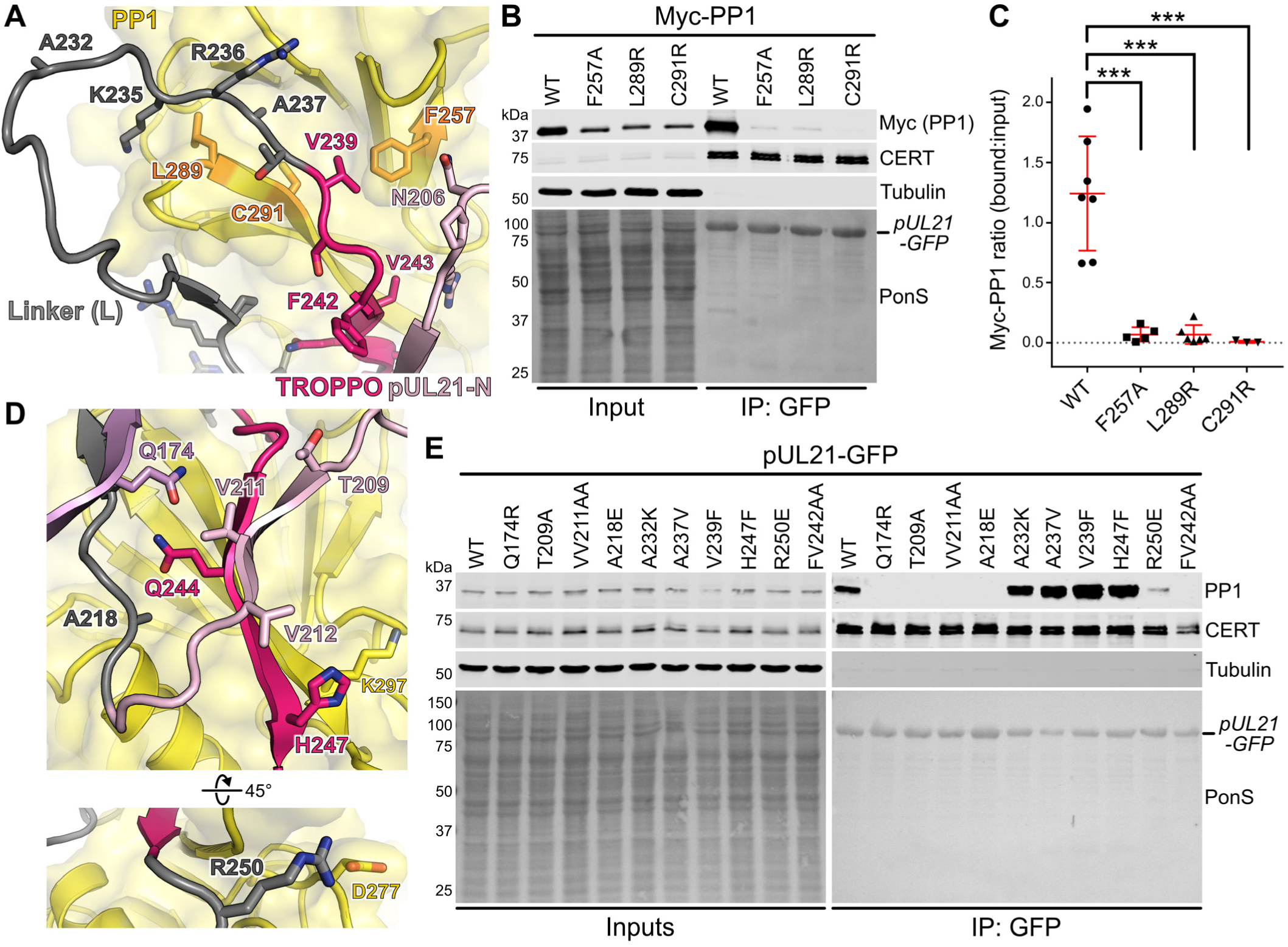
Site-directed mutagenesis supports pUL21 binding the hydrophobic RVxF groove of PP1. **A** Model of the pUL21:PP1 complex, highlighting predicted interactions with PP1 residues important for binding regulatory subunits containing RVxF motifs. Structures are shown as ribbons for PP1 (yellow), the pUL21 linker (grey), TROPPO (bright pink), and residues of pUL21-N missing from the crystal structure (light pink) (18). Selected residues shown as sticks. **B** HEK293T cells were co-transfected with plasmids expressing pUL21-GFP and wild-type (WT) or mutated Myc-tagged human PP1α. At 24 hours post-transfection the cells were lysed, subjected to immunoprecipitation using a GFP affinity resin, and captured proteins were subjected to SDS-PAGE and immunoblotting using the listed antibodies. Ponceau S (PonS) staining of the nitrocellulose membrane before blocking is shown, confirming efficient capture of GFP-tagged proteins. Figure is representative of at least three independent experiments (see below). **C** Quantitation of Myc-PP1 co-immunoprecipitation by pUL21-GFP. Ratio of bound to input signals as determined by densitometry is shown as individual points and mean ± SD from seven (WT), five (F257A and L289R) or three (C291R) independent experiments. One-way ANOVA with Holm-Sidak’s multiple comparisons test was used for the statistical analysis (***p < 0.001). **D** Model of the pUL21:PP1 complex, highlighting pUL21 residues at the interfaces with PP1 and pUL21-N. Ribbons are shown with selected residues represented as sticks, coloured as in (**A**). **E** HEK293T cells were transfected with plasmids expressing wild-type (WT) or mutated HSV-1 pUL21-GFP. At 24 hours post-transfection the cells were lysed, subjected to immunoprecipitation using a GFP affinity resin, and captured proteins were subjected to SDS-PAGE and immunoblotting using the listed antibodies. Ponceau S (PonS) staining of the nitrocellulose membrane before blocking is shown, confirming efficient capture of GFP-tagged proteins. Figure is representative of two independent experiments.

For pUL21, a panel of mutations were designed to probe the predicted interactions of residues at the interface with both PP1 and pUL21-N. Five mutations were designed to disrupt the interaction: Q174R, which would replace a buried residue on pUL21-N sheet β14 that contacts Q244 of the TROPPO motif (Fig. 4D); T209R, which would replace a small buried side chain that contacts pUL21-N with a large charged side chain (Fig. 4D); VV211AA, where two hydrophobic side chains (including one that is buried) in the sheet that bridges the TROPPO motif and pUL21-N β1 are removed (Fig. 4D); A218E, where a small hydrophobic residue that interacts with a surface hydrophobic patch on PP1 is replaced with a charged residue (Fig. 4D); and R250E, where the charge of a side chain that interacts with the PP1 D277 side chain is inverted (Fig. 4D). Three mutations were designed to enhance PP1 binding: A237V, substituting alanine for the canonical RVxF valine residue (Fig. 4A); V239F, substituting valine for the canonical RVxF phenylalanine residue (Fig. 4A); and H247F, substituting for the equivalent residue in GADD34 where the larger hydrophobic side chain could make more extensive interactions with the aryl portion of the PP1 K297 side chain (Fig. 4D). A control mutation A232K was also introduced, as substitution of a small hydrophobic side chain for a large charged one on a mobile surface-exposed loop (Fig. 4A) would be predicted to leave PP1 binding unaltered.

HEK293T cells were transfected with WT or mutant pUL21-GFP and the coIP of endogenous PP1α was monitored by immunoblotting (Fig. 4E). Pleasingly, all designed mutations had the predicted effect: PP1α coIP was abolished by the Q174R, T209A, VV211AA and A218E substitutions, in addition to the control FV242AA mutation, and the R250E substitutions dramatically reduced binding. All proteins were present at similar abundance in the input samples and all retained binding to CERT, confirming their correct folding. As predicted, the surface-loop substitution A232K did not dramatically alter PP1α coIP. The A237V, V239F and H247F mutants all co-precipitated more PP1α than did WT pUL21-GFP, suggesting that all three have higher affinity for the PP1 catalytic subunit. Taken together, these mutagenesis results strongly support the predicted structure of the pUL21:PP1 complex.

### Alphaherpesvirus proteins compete directly with cellular regulatory proteins for PP1 binding

Site-directed mutagenesis confirmed the predictive power of the pUL21:PP1 complex, but the affinity of the interaction between PP1 and alphaherpesvirus pUL21 homologues remained unknown. Attempts to perform ITC using recombinant full-length pUL21 and PP1 catalytic domain were unsuccessful as neither protein could be concentrated sufficiently to serve as titrant in the syringe. Similarly, attempts to purify full-length pORF38 following recombinant expression in *E. coli* were frustrated by low protein expression and solubility. As the R region of the linker and C-terminal domain are dispensable for PP1 binding, we designed expression constructs encoding the pUL21 and pORF38 N-terminal domains plus the Linker region up to the end of the TROPPO motif (the NLT region) with a C-terminal hexahistidine tag (Fig. 5A). The pORF38-NLT construct was readily expressed and purified to concentrations suitable for ITC (Fig. 5B). ITC titration of pORF38-NLT into PP1 showed that the proteins form a 1:1 complex with affinity of 497 ± 143 nM (mean ± SEM, three independent experiments) (Fig. 5C and Table S1). This is substantially weaker binding than observed for the GADD34 peptide binding PP1 via fluorescence anisotropy (Fig. 1A) or ITC (29).

**Figure 5.**
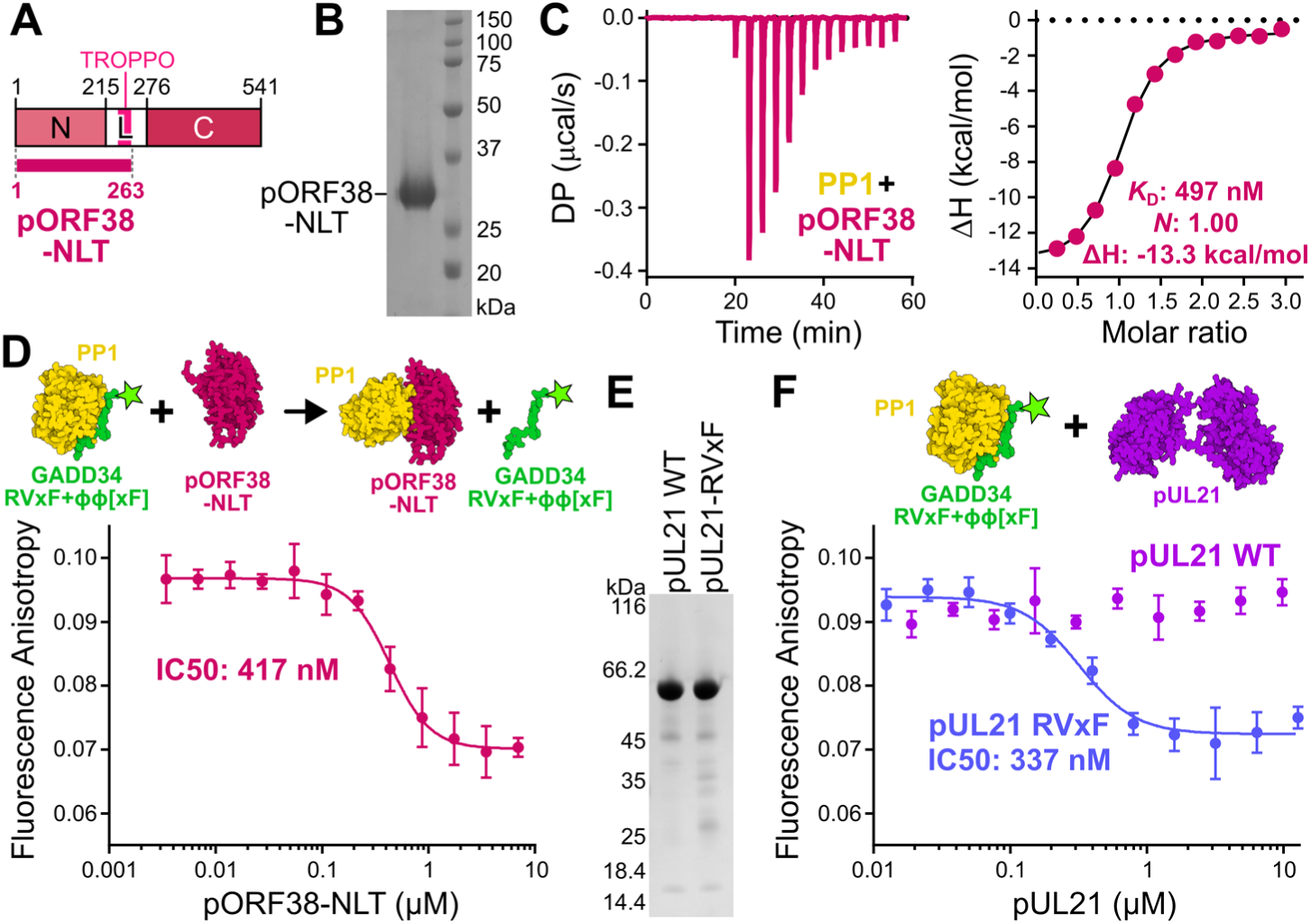
Alphaherpesvirus pUL21 homologues compete with cellular RVxF motifs for binding PP1. **A** Schematic of pORF38-NLT, which encodes the N-terminal domain plus Linker region to the end of the TROPPO motif. **B** Coomassie-stained SDS-PAGE of purified C-terminally hexahistidine-tagged pORF38-NLT. **C** ITC of PP1γ catalytic domain with pORF38-NLT. Baseline-corrected differential power (DP) versus time (left) and integrated change in enthalpy (ΔH) versus molar ratio (right) is shown. Figure is representative of three independent experiments (Table S1), and average values for affinity (*K*_D_), stoichiometry (*N*) and ΔH are shown. **D** Competition fluorescence anisotropy confirms that pORF38-NLT competes with RVxF+ϕϕ[xF] peptides for PP1 binding. A pre-formed complex of 2 nM fluorescent GADD34 RVxF+ϕϕ[xF] peptide with 250 nM PP1 was incubated with increasing concentrations of pORF38-NLT, resulting in displacement of the GADD34 peptide. Mean ± SD is shown for technical triplicate measurements. Average IC50 from three independent experiments is shown. **E** Coomassie-stained SDS-PAGE of purified C-terminally hexahistidine-tagged pUL21 WT and A237V+V239F (RVxF) mutant. **F** Competition fluorescence anisotropy of WT (purple) and RVxF (blue) pUL21. A pre-formed complex of fluorescent 2 nM GADD34 peptide with 250 nM PP1 was incubated with increasing concentrations of pUL21. Mean ± SD is shown for technical triplicate measurements. Average IC50 from three independent experiments is shown.

To definitively test whether pORF38 binds the same hydrophobic groove as cellular RVxF motifs, competition fluorescence anisotropy was performed via titration of pORF38-NLT into a pre-formed complex of PP1 plus the fluorescent GADD34 peptide. The liberation of GADD34 peptide at increasing concentrations of pORF38-NLT, as evidenced by a decrease in fluorescence anisotropy, confirmed that pORF38 competes with the RVxF peptide for PP1 binding (Fig. 5D), with an IC50 of 417 ± 11 nM (mean ± SEM, three independent experiments each performed in technical triplicate).

Similar competition fluorescence anisotropy experiments with purified full-length pUL21 (Fig. 5E) were performed in buffer optimised to enhance solubility of the protein (Tris pH 8.5 in place of HEPES pH 8). Under these conditions PP1 binds the GADD34 peptide with slightly higher affinity (55 ± 21 nM, mean ± SEM of three independent experiments performed in technical triplicate; Fig. S2). Full-length pUL21 is unable to out-compete GADD34 for binding to PP1 (Fig. 5F), suggesting that the affinity of PP1 for full-length pUL21 is lower than for pORF38 – consistent with the observation that PP1α is consistently co-immunoprecipitated more efficiently by pORF38-GFP than by pUL21-GFP (Fig. 2B and Fig. 3H of (20)). To test whether optimisation of the TROPPO sequence to better match a canonical RVxF motif could enhance binding, we generated, expressed and purified a pUL21-H_6_ mutant where residues A237 and V239 had been substituted for valine and phenylalanine, respectively (pUL21-RVxF, Fig. 5E). Fluorescence anisotropy confirms that pUL21-RVxF competes with the GADD34 peptide for binding to the PP1 hydrophobic groove with an IC50 of 337 ± 31 nM (mean ± SEM, three independent experiments each performed in technical triplicate).

## Discussion

pUL21 is required for efficient replication and spread of HSV-1 (20, 34–36). We previously identified HSV-1 pUL21 as a PP1 regulatory protein and that it contained a novel SLiM required for PP1 binding, the TROPPO motif (20). The absence of identifiable sequence homology with other known PP1-associated SLiMs (10) raised the tantalising possibility that the TROPPO motif binds PP1 via a novel surface that could be targeted for the development of antiviral compounds. Combining structure prediction, site-directed mutagenesis, immunoprecipitation and biophysical characterisation, we now show that the TROPPO motif binds the same hydrophobic surface of PP1 as cellular RVxF and ϕϕ[xF] SLiMs. Competition fluorescence anisotropy confirms that TROPPO-containing proteins compete directly with cellular RVxF and ϕϕ[xF] motifs for binding to PP1 (Fig. 5).

PP1 catalytic domains are unlikely to exist in their *apo* form within cells (10) and, as such, pUL21 and pORF38 must compete with cellular regulatory proteins to access PP1 in infected cells. There are over 200 mammalian PP1 regulatory proteins (9, 12) and their affinities for PP1 range from ∼10 nM (15, 37) to ∼1 µM (38). Given the competition for binding, it is puzzling that pORF38 has evolved to bind PP1 with only modest affinity (∼500 nM) (Fig. 5C). The PP1-binding of pUL21 must be even lower, as unlike pOPF38-NLT wild-type pUL21 cannot displace a GADD34 RVxF- and ϕϕ[xF]-containing peptide from PP1. One striking feature of the predicted pUL21:PP1 complex is that the TROPPO motif containing the ϕϕ[xF] motif lies sandwiched between PP1 and the pUL21-N, forming part of an extended β-sheet that bridges core β-sheets across the two proteins (Fig. 2A). While the structure of regulatory protein spinophilin (15) showed a sheet-turn-sheet binding PP1 at the ϕϕ[xF] interacting region, adding two strands to the PP1 core β-sheet, we are unaware of any examples where the RVxF and ϕϕ[xF] form a structured bridge between PP1 and another folded domain. The formation of this extended sheet, with concomitant burial of hydrophobic residues like F242 into a newly-extended hydrophobic core of pUL21-N (Fig. 3H), would stabilise the interaction between pUL21 and PP1 once formed. Such cooperativity would restrain the TROPPO motif in the correct conformation for PP1 binding despite the modest affinity (39).

Published RVxF consensus sequences (13, 40, 41) invariably have a valine or isoleucine and phenylalanine or tryptophan in the second and fourth position, respectively, whereas in pUL21 and pORF38 the equivalent residues are alanine and valine (Fig. 3). It is not clear that there are structural constraints preventing the virus from evolving an optimal RVxF motif since, like most regulatory protein regions that bind PP1 (12), the linker region of pUL21 is intrinsically disordered (20). Mutation of A237 to valine or V239 to phenylalanine enhances the coIP of PP1α by pUL21 in transfected cells (Fig. 4E) and the A237V+V239R double substitution gives pUL21-RVxF the ability to compete with the GADD34 peptide for PP1 binding (Fig. 5F). HSV-1 encodes a second PP1 regulatory protein, ICP34.5, that has high sequence homology to GADD34 and binds PP1 via a canonical RVxF motif (17, 42). There is thus nothing to prevent HSV-1 encoding a protein with a *bona fide* RVxF motif, and we hypothesise that selective pressure related to the molecular function of pUL21 has driven it to retain a suboptimal RVxF motif. Given the high degree of TROPPO sequence conservation across pUL21 homologues and corresponding absence of canonical RVxF motifs, we conclude that this evolutionary constraint must be common to all alphaherpesviruses.

What might drive these viruses to maintain a low-affinity PP1 recruitment domain? One clue comes from *in vitro* evolution experiments, where serial passage of HSV-1 encoding pUL21 mutated to prevent PP1 binding led to compensatory mutations in the virus-encoded kinase pUS3 (20). These two proteins have overlapping substrates, with pUL21-mediated dephosphorylation directly antagonising pUS3 kinase activity. A balance of activity is clearly critical for the virus: abolishing expression of either pUS3 or pUL21 severely diminishes virus replication in cultured keratinocytes, as does removal of pUL21 PP1 binding, but passage of the PP1-binding pUL21 mutants led to rapid accumulation of mutations that lowered pUS3 kinase activity and restored virus fitness. The peak of pUL21 expression is at late times (18 hours) post-infection in cultured keratinocytes, whereas pUS3 abundance peaks at 6 hours post-infection and then declines slightly (43). We hypothesise that the PP1-binding motif of pUL21 is specifically tuned to prevent effective PP1 binding at early times post-infection, when pUS3 activity is presumably pro-viral. Instead, the presence of a suboptimal PP1 interaction motif would allow pUL21 to effectively compete for PP1 binding only at late times post infection and in specific subcellular locations, for example the nuclear envelope, where the protein is highly abundant (20). This fine-tuning of activity would be especially important for a multi-functional protein like pUL21, with roles cell-to-cell spread (20, 34, 36, 44–46), preventing futile genome release by nascent viral capsids (35, 47) and virus assembly (45, 46, 48, 49), in addition to its role in regulating viral nuclear egress (44, 45, 50–52). High affinity binding to PP1 could recruit phosphatase activity to the wrong cellular or viral complexes, or it could inappropriately sequester PP1 and thereby prevent other PP1-mediated dephosphorylation events within the cell that promote virus replication. Similar fine-tuning of PP1 recruitment has been observed for HIV, where mutation of the suboptimal QVCF motif to RVCF enhanced PP1 binding but reduced the ability of Tat to stimulate HIV transcription (5).

In conclusion, we show that the alphaherpesvirus pUL21 homologues bind the same PP1 hydrophobic surface groove as cellular RVxF and ϕϕ[xF] motifs. Extensive site-directed mutagenesis supports a predicted structure of the pUL21:PP1 complex where intrinsically disordered regions of the pUL21 linker fold into an extended β-sheet that bridges the cores of PP1 and the pUL21 N-terminal domain. The conservation of suboptimal binding residues within the region structurally equivalent to the cellular RVxF implies that the viruses have deliberately regulated PP1 binding affinity to support optimal virus replication. This highlights the careful regulation of kinase and phosphatase activity required to support efficient virus replication and spread. Our work also highlights how PP1 can be recruited by proteins lacking any identifiable PP1-binding SLiM, suggesting that the pool of viral and cellular PP1 regulatory proteins may be larger than previously imagined.

## Acknowledgements

We thank Owen Tutt (University of Cambridge) for technical assistance. For the purpose of open access, the authors have applied a Creative Commons Attribution (CC BY) licence to any Author Accepted Manuscript version arising from this submission.

## Data availability

The models of pUL21 and pORF38 in complex with PP1 have been deposited to the University of Cambridge Data Repository (DOI). The authors confirm that all other data supporting the findings of this study are available within the article and/or its supplementary materials.

---

**Figure S1.**
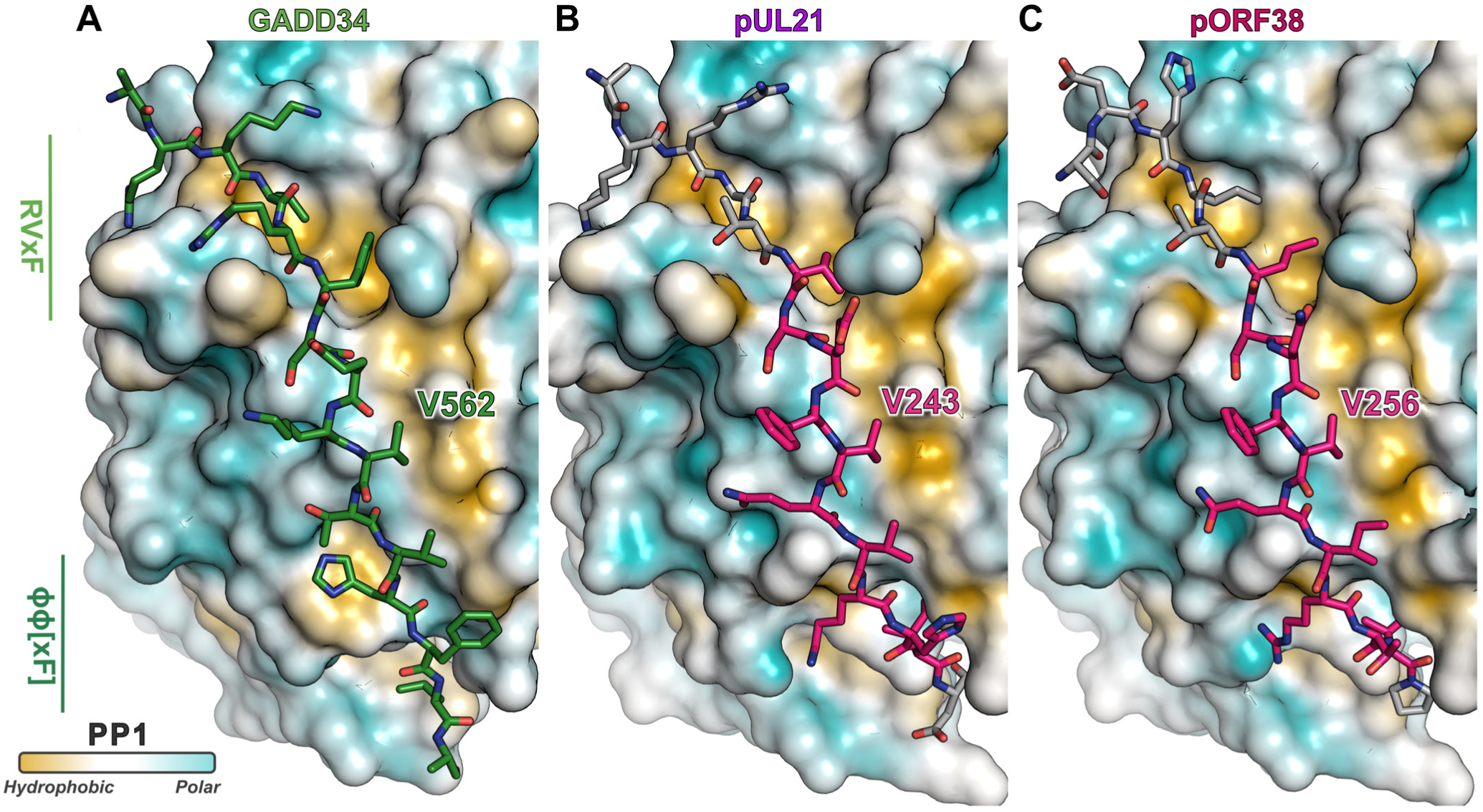
Comparison of (predicted) peptide conformations at the PP1 hydrophobic groove. **A–C** PP1 is shown as a molecular surface coloured by mean lipophilicity, from orange (hydrophobic) to cyan (polar). **A** Structure of the crystal structure of PP1α catalytic domain in complex with the GADD34 RVxF+ϕϕ[xF] motif (PDB 4XPN) (29). The GADD34 peptide is shown as sticks with green carbon atoms. **B** AlphaFold2-Multimer model of PP1γ catalytic domain in complex with HSV-1 pUL21. The TROPPO motif (pink carbon atoms) and flanking residues (grey alpha carbons) are shown as sticks. **C** AlphaFold2-Multimer model of PP1γ catalytic domain in complex with VZV pORF38. The TROPPO motif (pink carbon atoms) and flanking residues (grey alpha carbons) are shown as sticks.

**Figure S2.**
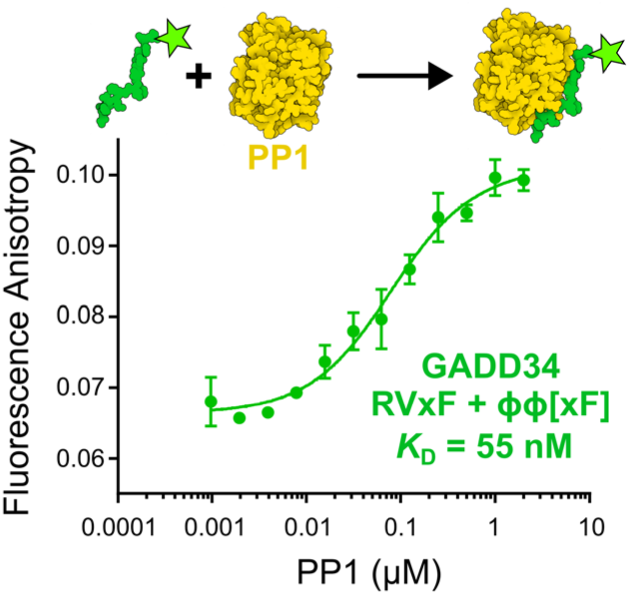
Association of GADD34 peptide and PP1 in Tris pH 8.5 buffer. Fluorescence anisotropy of PP1γ catalytic domain (residues 7–300) binding 2 nM fluorescently labelled peptide containing the GADD34 RVxF and ϕϕ[xF] motifs (residues 552–568). Mean ± SD is shown for technical triplicate measurements. Affinity (*K*_D_) is average of three independent experiments.

