## Supplementary material for "Alphaherpesvirus pUL21 homologues use non-canonical motifs to compete with cellular adaptors for protein phosphatase 1 binding": Table S1

**Table S1. Isothermal titration calorimetry (ITC) of PP1 with TROPPO-containing peptides and proteins.** Data for independent experiments are shown. For all, the cell contained C-terminally $His_6$-tagged mouse PP1γ (7–300). –, no binding detected.

| Titrant | [Titrant] in syringe (μM) | [PP1] in cell (μM) | $K_D$ (nM) | ΔH (kcal/mol) | ΔG (kcal/mol) | -TΔS (kcal/mol) | N (sites) |
|---|---|---|---|---|---|---|---|
| pUL21-TROPPO (234–250) | 1000 | 58.8 | – | – | – | – | – |
| | 1000 | 54.5 | – | – | – | – | – |
| pORF38-TROPPO (246–263) | 1000 | 79.6 | – | – | – | – | – |
| | 500 | 17.0 | – | – | – | – | – |
| | 500 | 16.0 | – | – | – | – | – |
| pORF38-NLT-$His_6$ (1–263) | 240 | 17.0 | 719 | -13.1 | -8.38 | 4.76 | 1.21 |
| | 147 | 10.0 | 540 | -13.3 | -8.55 | 4.72 | 0.965 |
| | 153 | 9.5 | 231 | -13.6 | -9.06 | 4.5 | 0.838 |
